# A curated collection of transcriptome datasets to investigate the molecular mechanisms of immunoglobulin E-mediated atopic diseases

**DOI:** 10.1101/525477

**Authors:** Susie S. Y. Huang, Fatima Al Ali, Sabri Boughorbel, Mohammed Toufiq, Damien Chaussabel, Mathieu Garand

## Abstract

Prevalence of allergies has reached ~50% of industrialized populations and with children under ten being the most susceptible. However, the combination of the complexity of atopic allergy susceptibility/development and environmental factors has made identification of gene biomarkers challenging. The amount of publicly accessible transcriptomic data presents an unprecedented opportunity for mechanistic discoveries and validation of complex disease signatures across studies. However, this necessitates structured methodologies and visual tools for the interpretation of results. Here, we present a curated collection of transcriptomic datasets relevant to immunoglobin E (IgE)-mediated atopic diseases (ranging from allergies to primary immunodeficiencies). 30 datasets from the Gene Expression Omnibus (GEO), encompassing 1761 transcriptome profiles, were made available on the Gene Expression Browser (GXB), an online and open-source web application that allows for the query, visualization, and annotation of metadata. The thematic compositions, disease categories, sample number, and platforms of the collection are described. Ranked gene lists and sample grouping are used to facilitate data visualization/interpretation and are available online via GXB (http://ige.gxbsidra.org/dm3/geneBrowser/list). Dataset validation using associated publications showed good concordance in GXB gene expression trend and fold-change.

Database URL: http://ige.gxbsidra.org/dm3/geneBrowser/list

## INTRODUCTION

Allergic disease is highly prevalent and currently reaches ~50% of the populations in industrialized nations (1). Although the generation of allergic responses is well-understood, the early sensitization steps and factors contributing to the development of immunoglobin E (IgE)-mediated diseases remain unclear. IgE is the major mediator of atopic response in humans, whereas atopy represents the predisposition to become over IgE-sensitized to allergens. However, not all encounters with a potential allergen will lead to sensitization. Similarly, not all sensitizations will result in a symptomatic allergic response even in atopic individuals.

The effect of IgE spans across multiple systems. In the circulation system, IgE increases flow and permeability of the blood vessels, fluid and protein in tissues, as well as flow to the lymph nodes. In the airway, IgE decreases air conduct diameter, increases mucus congestion, and can induce blockage. In the gastro-intestinal (GI) tract, IgE increases fluid secretion, peristalsis, and expulsion-diarrhea. The immunoglobin has also been suggested to play a role in the defense against parasite infection and as a general gate keeper for any foreign materials entering the body (2,3). Ultimately, IgE is involved in the normal spectrum of reaction to expulse foreign material from the body; hence, protecting by elimination. Nevertheless, over sensitization, which is developed by unknown mechanism(s), can lead to imbalance and pathology in affected individuals.

Two main groups of immune signals initiate the production of IgE in response to an antigen: 1) the signals that drive the differentiation of CD4 naive T cell to T helper type 2 (Th2) cells and 2) the cytokines and co-stimulatory molecules secreted by Th2 cells, which subsequently promote T follicular helper cells-induced immunoglobulin B cells class switch towards IgE production. Antigen characteristics, such as concentration and localization of the encounter (i.e. tissue, mucosa, circulation), can also affect Th2 cell induction. IL-4, IL-13, and STAT6 are key mediators of Th2 responses and IgM class switch to IgE. IL-4 secretion and mast cell CD40 surface expression also contribute to the IgE class switch to IgE at the site of allergic reaction.

Systemic levels of IgE alone is not a sufficient indicator for allergy risk (4). Peripheral blood IgE level can increase upon sensitization, but not reliable enough for deducing a diagnosis of allergy or allergen type (5). Concentration, binding strength or affinity, specificity, and portion of specific IgE to total IgE are all factors in translating a humoral IgE response into a clinical symptom (6). However, genetic component exists in allergic disease. Studies have demonstrated strong heritable components of allergic diseases and atopy, estimated at 33%-76% (7,8). Genetic components of food allergy and asthma have also been reviewed (9–12). Although single susceptibility gene have been identified for certain allergies (13–16), most genome-wide association studies reported multiple loci of susceptibility or gene regions to a wide range of allergy development (17–19). Environmental factors further complicate allergic susceptibility - the hygiene hypothesis of allergic susceptibility has received much attention in the last few decades (20). Other exposure-related factors (e.g. diet, pollution, tobacco smoke) are also likely to have contributed to the increasing susceptibility to allergic diseases in developed nations (1).

IgE may also be implicated in some primary immunodeficiency diseases (PIDs). For CD40 ligand and CD40 deficiency, circulating IgE+ plasma cells are absent (21). For others, IgE plasma levels are elevated: DOCK8 deficiency, AD-HIES Job’s syndrome, Comel-Netherton syndrome, PGM3 deficiency, IPEX (immune dysregulation, polyendocrinopathy, enteropathy X-linked), and Tyk2 deficiency. Large phenotypic heterogeneity is observed among these PIDs; however, the underlying mechanisms is not completely understood. This knowledge gap mirrors our limited understanding of the involved gene products mediating phenotypes atopic diseases.

There is a crucial need to pursue genetic markers of an atopic constitution amidst growing concern over the increasing abundance of IgE atopy and that current treatments remain mostly palliative. As both environmental interaction and genetic predisposition forms the determinant of allergy development, extrapolating potential gene markers of allergy atopy from a broad range of studies with associated clinical data is highly relevant, particularly given the high variations of the disease clinical phenotypes. With the advances in high through-put platforms for transcriptomic analyses, a large number of gene expression datasets is regularly deposited on publicly accessible repositories such as the NCBI Gene Expression Omnibus (GEO). GEO currently hosts over 90 000 datasets comprising over 2 million samples (https://www.ncbi.nlm.nih.gov/geo/) and presents an enormous opportunity for data mining across multiple studies.

The web-based Gene Expression Browser (GXB) is an open-source interface that allows for custom compilation of selected datasets (i.e. of interest to users) and facilitates visualization of gene expression data. GXB has been previously described (22) and used to generate a number of Data Notes (23–28). As well, GXB is a useful tool for novel gene/function discovery (29) and in system re-analysis approaches (30).

A search strategy was implemented to identify GEO datasets relevant to IgE-related atopic disease, including PIDs, and uploaded those datasets to the GXB platform online. The associated study metadata, such as the detailed gene information, relevant literature, study design, and sample information, were also uploaded to facilitate in-depth interpretation. In creating this collection of datasets, we aimed provide a resource which will facilitate risk prediction and interception of allergic diseases. The datasets were retrieved from publicly available GEO series and selected by relevance to IgE and atopic diseases and filtered by analysis platform and species. As previously demonstrated, data mining and re-analysis/re-interpretation of large and publicly available dataset is a promising avenue (31) to elucidate complicated diseases such as IgE-related atopy.

## METHODS

### Justification of data selection and filter

The focus of the GEO dataset selection was primarily on whether or not the dataset involved 1) IgE production or function or 2) if the pathology studied implicated directly or indirectly IgE production or function (Table 1A). We ensured a broad search approach by combining the 7 independent search results (Table 1A) using the “Merge collection” function available on “My NCBI” (https://www.ncbi.nlm.nih.gov/myncbi/) (Table 1B) and by performing another search using an assembly of all the terms used in Table 1A (Table 1C). 203 potentially relevant datasets were identified from the initial query (Table 1D). The query results were then manually filtered to restrict datasets to human sample, expression profiling by microarrays, and relevance to IgE-related atopic diseases. The process involved inspecting the study description, design, and sample type for each dataset and resulted in 30 curated datasets. The following criteria were deemed very important: clear description of tissue type, comparisons between patient vs healthy or stimulated vs unstimulated samples, and disease category implicating IgE. Furthermore, datasets/studies that are indirectly relevant to IgE-mediated atopic disease, such as gene expression in B cells after tonsillectomy, were also included as they were deemed valuable to 1) the discovery of putative novel gene-disease association, 2) to improving our knowledge of adaptive immunity, and/or 3) to increasing our knowledge about factors affecting IgE production.

**Table 1.** Search strategy for dataset retrieval from the Gene Expression Omnibus (GEO) database.

A. Search strategy applied for retrieval of relevant datasets from GEO.
| Description | Search strategy | # of dataset |
| --- | --- | --- |
| Dataset relevant to hypersensitivity, hyper-IgE syndrome, and IgE in syndromes, disorders, diseases, and the regulation/phenotype in general of IgE. | ("Hypersensitivity" OR "hyper ige syndrome" OR HIES OR hyper-IgE OR IgE OR "immunoglobulin e") AND (syndrome OR syndromes OR disorder OR disorders OR "job's syndrome" OR disease OR diseases OR regulation OR modulation OR change OR changes OR elevated OR decreased OR deficiency OR phenotype) AND "gse"[Filter] | 130 |
| Dataset relevant to hyper-IgE and IgE. | ("hyper ige syndrome"[All Fields] OR HIES[All Fields] OR hyper-IgE[All Fields] OR IgE[All Fields] OR "immunoglobulin e"[All Fields]) AND "gse"[Filter] = 75 | 75 |
| Dataset relevant to hyper-IgE and IgE in allergenic/antigenic reactions, syndromes, disorders, diseases, and the regulation/phenotype in general of IgE. | ("Hyperimmunoglobulinemia E syndrome" OR "hyper ige syndrome" OR HIES OR hyper-IgE OR IgE OR "immunoglobulin e") AND ("Allergens"[Mesh] OR "Antigens"[Mesh] OR syndrome OR syndromes OR disorder OR disorders OR "job's syndrome" OR disease OR diseases OR regulation OR modulation OR change OR elevated OR decreased OR deficiency OR phenotype) AND ("gse"[Filter]) | 62 |
| Dataset relevant to IgE receptor, hyper-IgE, IgE and mast cells in hypersensitivity and allergy. | (FcεR OR "IgE receptor" OR "hyper ige syndrome"[All Fields] OR HIES[All Fields] OR hyper-IgE[All Fields] OR IgE[All Fields] OR "immunoglobulin e"[All Fields] OR "mast cells"[All Fields] OR "Mast Cells"[Mesh]) AND ("hypersensitivity"[MeSH Terms] OR "allergy and immunology"[MeSH Terms] OR ("hypersensitivity"[MeSH Terms] OR "allergy and immunology"[MeSH Terms] OR ("hypersensitivity"[MeSH Terms] OR "allergy and immunology"[MeSH Terms] OR ("hypersensitivity"[MeSH Terms] OR "allergy and immunology"[MeSH Terms] OR allergy[All Fields])))) AND "gse"[Filter] | 48 |
| Dataset relevant to hyper-IgE and IgE in hypersensitivity and allergy. | ("hyper ige syndrome"[All Fields] OR HIES[All Fields] OR hyper-IgE[All Fields] OR IgE[All Fields] OR "immunoglobulin e"[All Fields]) AND ("hypersensitivity"[MeSH Terms] OR "allergy and immunology"[MeSH Terms] OR allergy[All Fields]) AND "gse"[Filter] | 33 |
| Dataset relevant to IgE in dermatitis, hypersensitivity, allergy, and mast cells. | ((("dermatitis"[MeSH Terms] OR dermatitis[All Fields]) OR ("hypersensitivity"[MeSH Terms] OR "allergy and immunology"[MeSH Terms] OR allergy[All Fields]) OR ("mast cells"[MeSH Terms] OR mast cells[All Fields])) AND (IgE[DESC] OR ("immunoglobulin e"[All Fields]))) AND "gse"[Filter] | 50 |
| Dataset relevant to hyper-IgE and IgE in immune system diseases and study about allergens. | ("hyper ige syndrome"[All Fields] OR HIES[All Fields] OR hyper-IgE[All Fields] OR IgE[All Fields] OR "immunoglobulin e"[All Fields]) AND "gse"[Filter] AND ("Immune System Diseases"[Mesh] OR "Allergens"[Mesh]) | 37 |

**B. Merged of the result from the search strategy described in A.**
| Description | Search strategy | # of dataset |
| --- | --- | --- |
| Merging the results of all 7 collections mentioned above. | Merge saved collections in "My NCBI"(https://www.ncbi.nlm.nih.gov/myncbi/) | 196 |
| ---> performed on human samples | Filtering | 143 |
| ---> performed on mouse samples | Filtering | 52 |

**C. Merged of the terms used in the search strategy described in A.**
| Description | Search strategy | # of dataset |
| --- | --- | --- |
| Merging all terms used in the 7 search strategy above. | (FcεR[All Fields] OR "IgE receptor"[All Fields] OR "mast cells"[All Fields] OR "Mast Cells"[Mesh] OR "hyper ige syndrome"[All Fields] OR HIES[All Fields] OR hyper-IgE[All Fields] OR IgE[All Fields] OR "immunoglobulin e"[All Fields] OR "Immunoglobulin E"[Mesh]) AND ("Immune System Diseases"[Mesh] OR dermatitis[All Fields] OR allergy[All Fields] OR ("Allergens"[Mesh] OR "Antigens"[Mesh]) syndromes[All Fields] OR disease[All Fields] OR "disorder"[All Fields] OR "job's syndrome"[All Fields] OR "regulation"[All Fields] OR modulation[All Fields] OR (change[All Fields] OR changes[All Fields]) OR elevated[All Fields] OR decreased[All Fields] OR deficiency[All Fields] OR phenotype[All Fields]) AND "gse"[Filter] | 117 |
| ---> performed on human samples | Filtering | 70 |
| ---> performed on mouse samples | Filtering | 37 |

**D. Final assembly and curation of search strategy**
| Description | Search strategy | # of dataset |
| --- | --- | --- |
| Merging of "B" and "C" results, only gene expression profiling by microarrays or high-throughput sequencing. | Assembly of the final list of dataset by merging collections B (196) and C (117) in "My NCBI"(https://www.ncbi.nlm.nih.gov/myncbi/) | 203 |
| ---> performed on human samples | Filtering | 115 |
| ---> performed on mouse samples | Filtering | 76 |
| **For human samples only:** |  |  |
| Manual inspection of dataset to confirm implication to IgE-related pathogenesis in human only. | Inspect all human dataset descriptions using the "GEO accession display tool" (https://www.ncbi.nlm.nih.gov/geo/query/acc.cgi) | 53 |
| Select gene expression by arrays only. | Filter platform type | 49 |
| Manual inspection of the type of study and design, the type of samples, and the disease groups. | Read full description and protocol of each dataset available on "GEO accession display tool" (https://www.ncbi.nlm.nih.gov/geo/query/acc.cgi) and inspect associated published article when necessary | 30 |

### The web-based GXB platform: a visualization tool for gene expression data

GXB is a valuable tool for training researchers about the reductionist investigative approaches (see description of such program in (31)). The creation of a web-based GXB platform was previously described in detail (22). In brief, the GXB is a simple interactive interface designed for visualization of large quantities of heterogenous data (Suppl. Figure 2). The platform allows for customizable data plots with overlapping metadata information, changeable sample order, as well as generation of sharable mini-URLs that encapsulate information about the display settings in use. The dataset navigation page allows for quick identification of datasets of interest either through filtering using pre-define lists or via query terms. The user has access to multiple functionalities within the GXB graphic interface. In brief, the data-viewing interface enables interactive browsing and graphic representation of large-scale data. This interpretable format displays ranked gene lists and expression results. The interface also allows for user flexibility in terms of changing how the gene list is ranked, the method used for ranking, sample grouping (i.e. disease type), sample sorting (i.e. gender or age) and view type (i.e. bar or chart). The end user can browse through the datasets, format graph for a selected gene within a dataset, and export data (i.e. annotation, FC, signal, groups). The original GEO data and annotation are accessible from the GXB interface (Downloads tabs) as well as from their GEO page (links provided on Study tab). The associated source code and instructions are publicly available (https://github.com/BenaroyaResearch/gxbrowser)], as well as the necessary R scripts (https://github.com/BenaroyaResearch/gxrscripts).

### Construction of the dataset collection on GXB

The selected datasets were downloaded from GEO in SOFT file format and uploaded (http://ige.gxbsidra.org/dm3/geneBrowser/list) via the GXB interface (accessed by navigating the Tools menu located in the top-right corner of the GXB webpage; Tools / Chips Loaded / Upload Expression Data). The SOFT files contain metadata and normalized signal intensity data, generated by methods indicated by the author(s). These SOFT files can be analyzed directly for differentially expressed genes; thus, no additional processing was required. Using the “Sample Set Annotation Tool” of the GXB interface, the datasets were annotated according to the information provided on GEO. The raw signal data type of the dataset (ex. raw signal, log2 transformed and etc.,) are also specified for each GEO entry. The default data display is in linear scale (see further details below in “Presentation of datasets”). When necessary, the sample annotation file (which is part of the SOFT file) were edited in order to add group information; as it was important to identify groups in order to compute fold change (FC). For each dataset, individual samples were grouped based on relevant study variables. Three datasets (GSE19190, GSE75603, and GSE8507) were split via GXB’s “Sample Set Annotation Tool/Group Sets” interface to provide a more meaningful group comparison (these datasets share the same GSE number in Table 2). Genes were then ranked based on FCs of the specified two-group comparison. All the information annotated and presented on GXB is assembled in a SQL database (Data Citation 1).

**Table 2.**
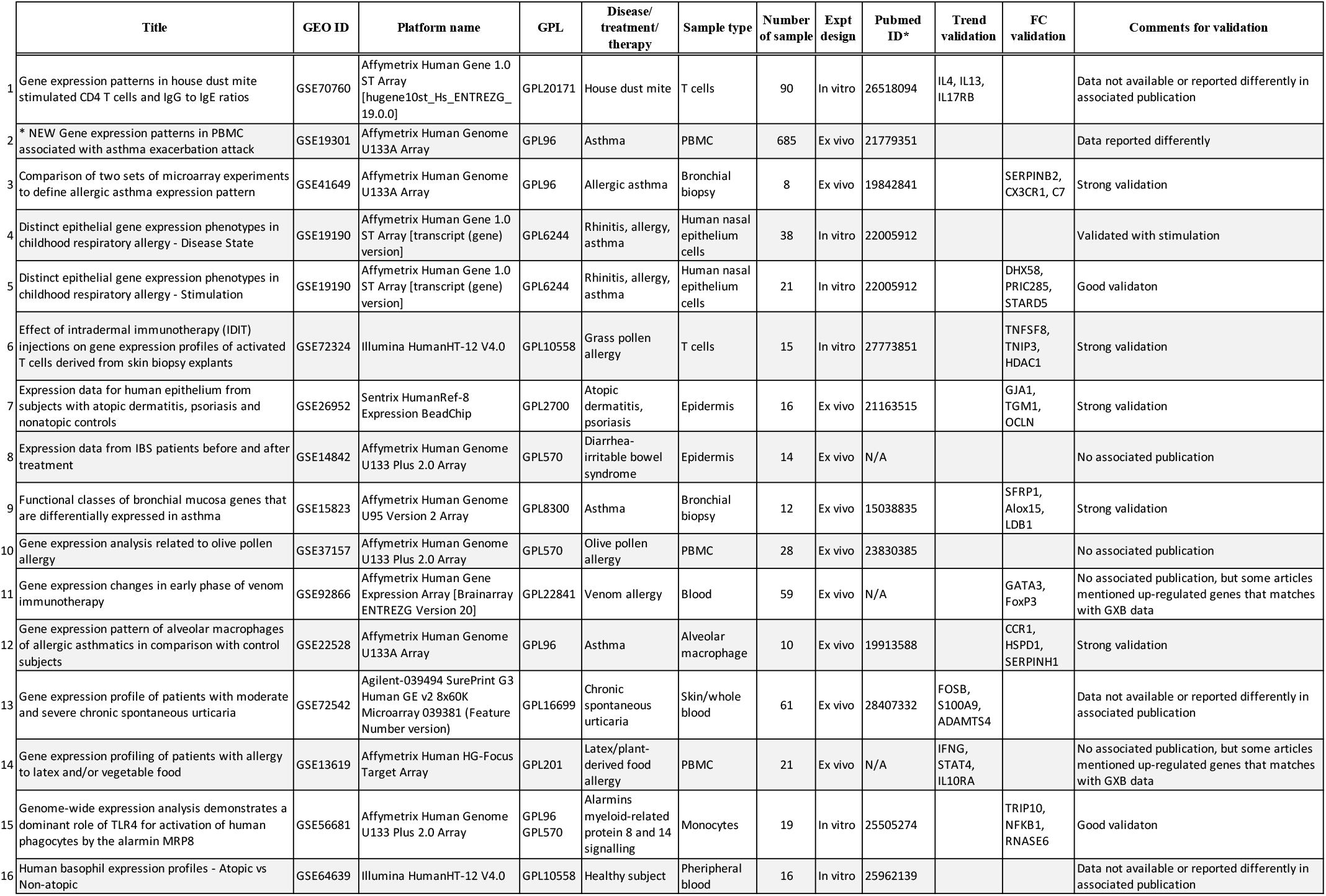
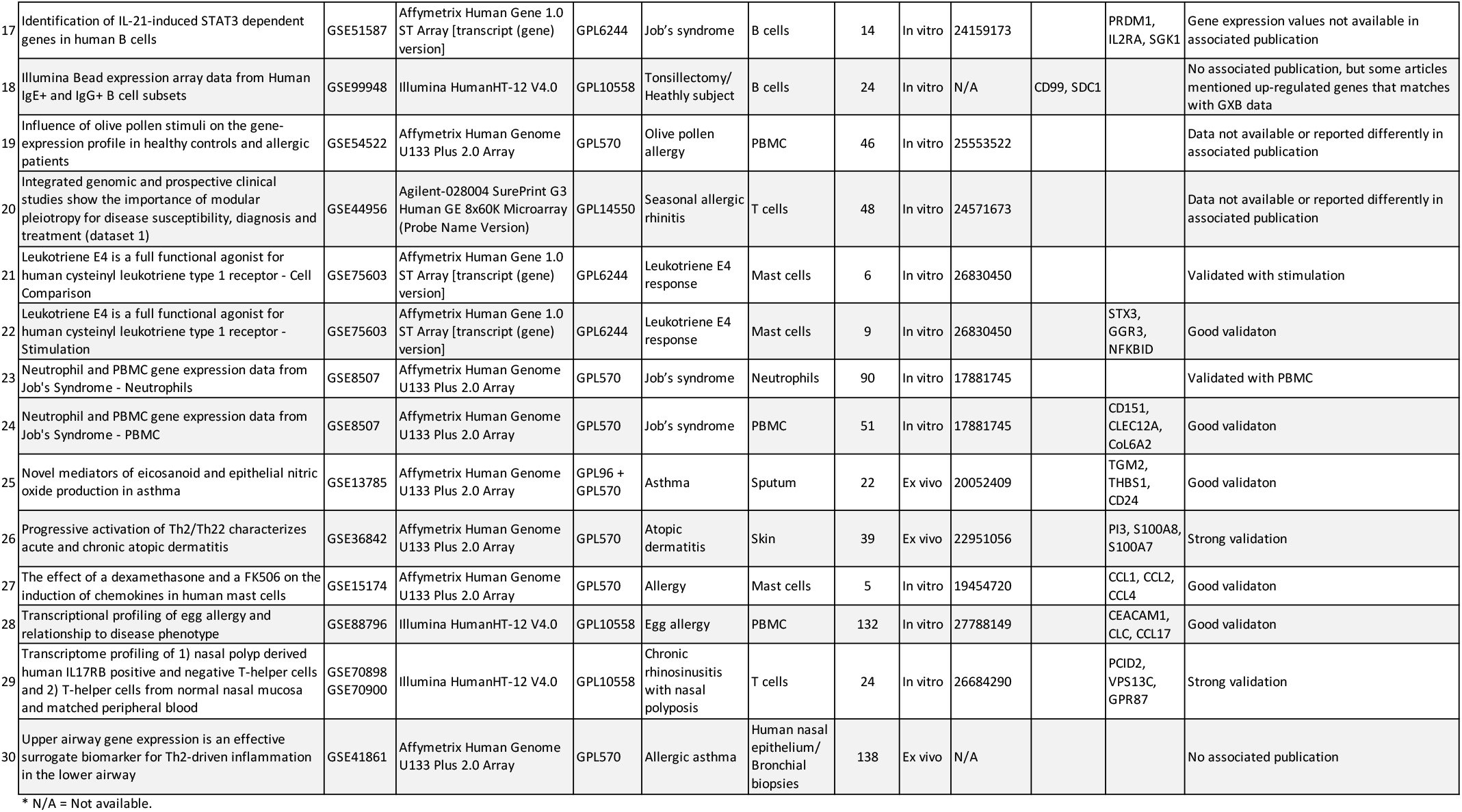
Descriptive summary of the dataset collection.

### Data availability

The curated datasets collected that have been described in this data note were assembled from the public repository NCBI GEO website: http://www.ncbi.nlm.nih.gov/gds/. In this study, we cited each dataset GEO accession number, and raw signal and annotation files are made available for download from the GXB web-application (http://ige.gxbsidra.org/dm3/geneBrowser/list).

## RESULTS

### Description of datasets

After applying the filtering strategy as previously described, we curated 30 datasets encompassing 1761 transcriptome profiles relevant to IgE-related atopic diseases. Detailed information on each dataset is presented in Table 2 and the summary of the data collection is presented in an aggregation of pie charts (Figure 1). The data collection includes a wide range of studies, sample types, as well as diseases. In total, 12 different microarray platforms are represented, with the majority being Affymetrix Human Array chips (various version). 17 datasets were generated from *in vitro* studies and 13 from *ex vivo* studies. Ten sample types are represented, with the most abundant being PBMC (n = 8) and airway epithelial cells (n = 5). Sample size of each dataset ranges from 5 to 628, with most studies having 10-50 samples. The dataset collection covered 7 main disease categories, including allergy, asthma, healthy responses, hyper IgE syndrome (HIES), dermatitis, atopic irritable bowel syndrome (IBS), and Urticaria. In the majority of the studies, comparisons are made between patients and health controls (n = 17), followed by stimulation (n = 6) and time course (n = 5). The frequency of the terms retrieved from the associated GEO descriptions and those derived from the MeSH/keywords of the published articles are visualized through word clouds presented in Figure 2.

**Figure 1.**
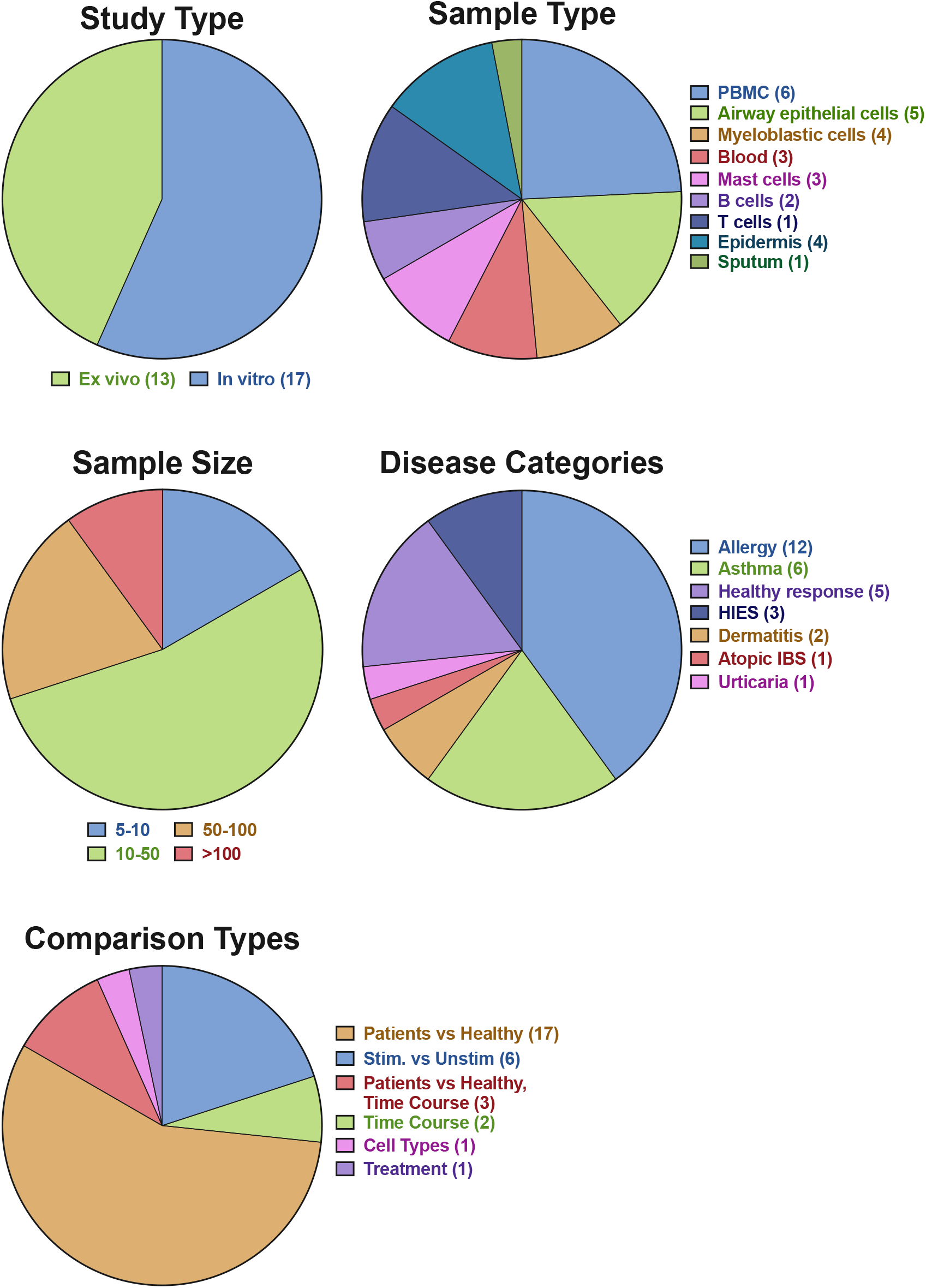
Characteristic of the curated dataset collection. Proportion and number of the study type, sample type, sample size, disease categories, and type of comparison performed for fold-change calculation on the GXB are depicted.

**Figure 2.**
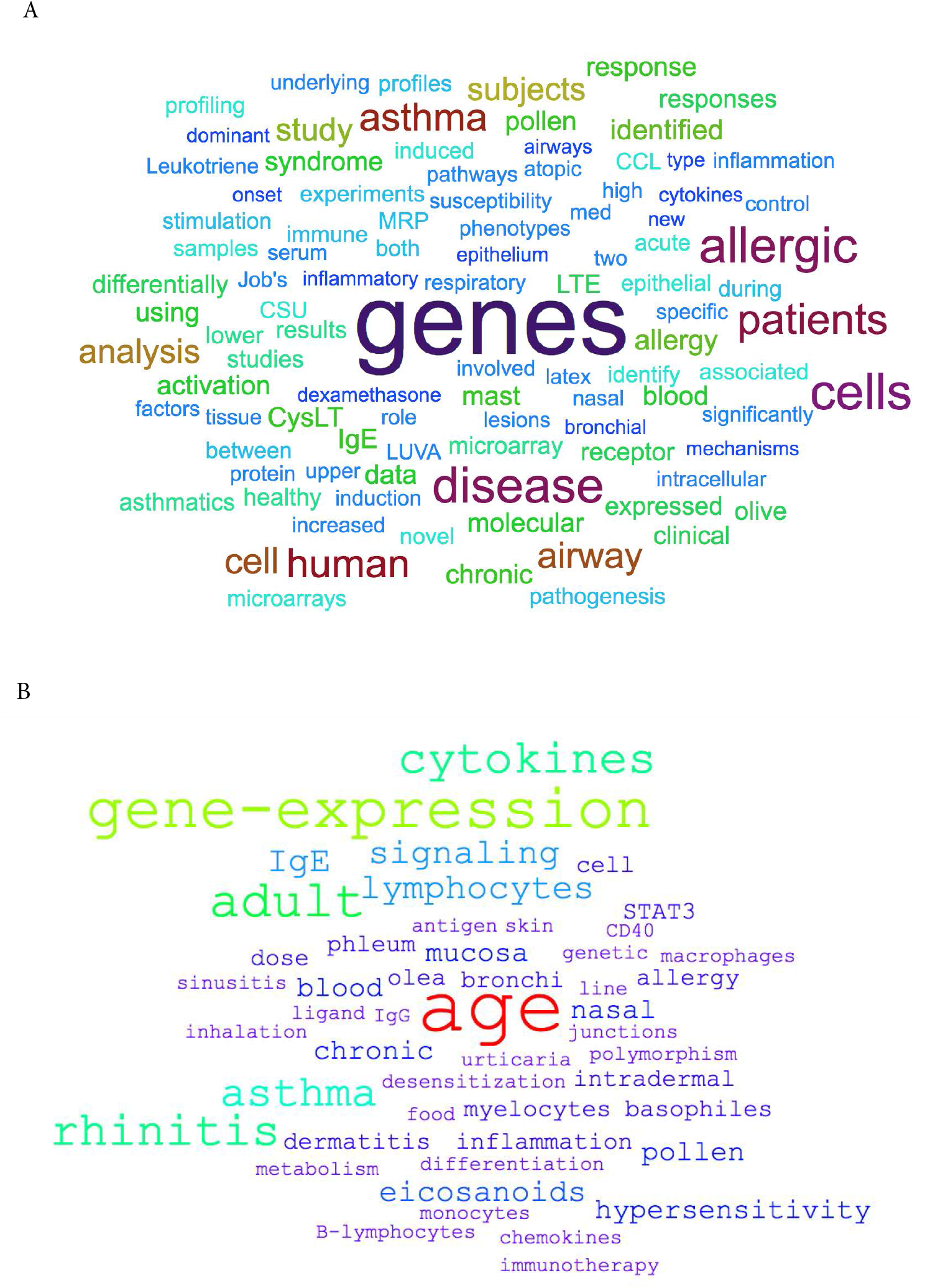
Word clouds depicting the frequency of (A) the terms retrieved from the associated GEO descriptions of the datasets and (B) those derived from the MeSH/keywords of the associated published articles.

### Presentation of datasets

On the graphic interface of GXB, genes expression values are displayed in linear scale. The original signal data type can be found under the Info tab of individual datasets. The associated metadata are available under the Samples tab and this information can be used for graphical overlaying (via “Overlays” dropdown menu). The FCs are also displayed in linear scale (see Supplemental Information for a detailed example of FC calculation). For FC analysis, each subject/sample are grouped according to the experimental design (experimental variable vs controls) and/or, if available, as per the corresponding publications. It is also possible to rank the genes based on expression difference by selecting “Advanced” in the “Rank Lists” dropdown menu of the graphic interface. This display option can be more robust than FC when low expression is observed in one group. It is important to note that integration and re-analysis of the datasets is not the intent of this collection. Therefore, a more meaningful use of this collection lies in the comparison of FC expression for a gene across multiple relevant datasets (i.e. reductionist approach (31).

### Dataset validation

Quality assessment of the datasets was performed by looking for key gene expression, i.e. marker genes, and highly differentiated genes as indicated in the associated publications. Table 2 includes the type of validation, the genes used, as well as the strength of the validation and the associated comments. Certain datasets were split in two to facilitate comparisons (GSE19190, GSE75603, and GSE8507); in these cases, validation is indicated for one of the two datasets as indicated in Table 2. Data validation was achieved for most datasets except for 6, due to having no publications (GSE14842, GSE37157, and GSE41861) or incomparable units (i.e. z-score) and/or absence of FC information (GSE64639, GSE54522 and GSE44956). Trend validations were done on 4 datasets that had no linked publications (GSE13619 and GSE99948) or had incomparable units to GXB (GSE70760 and GSE72542). But when possible, literature values were used for additional validation of the trend (i.e., GSE92688and GSE99948). Examples of the validation results are presented in Figure 3.

**Figure 3.**
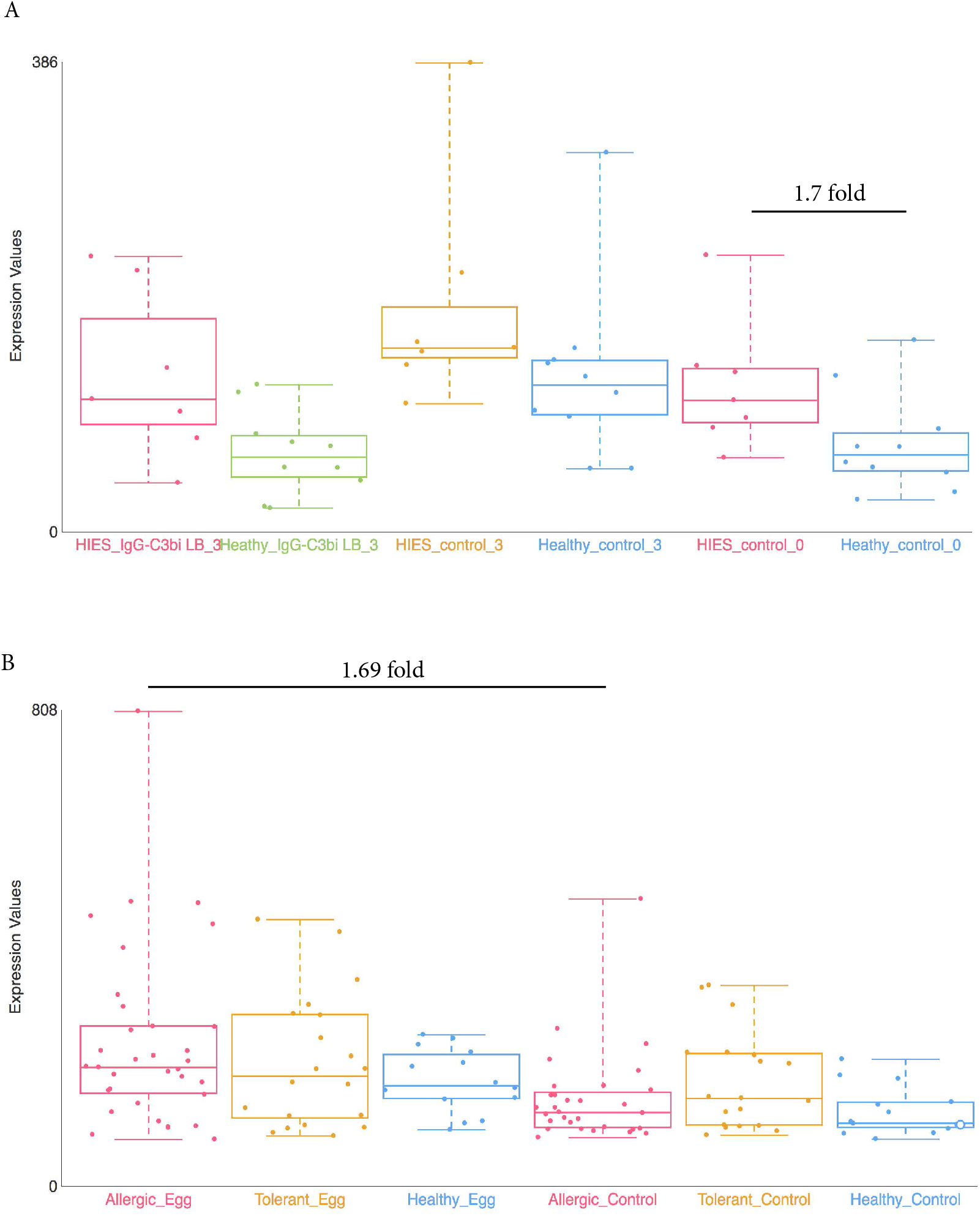
Examples of dataset validation. Differently expressed genes from 2 datasets were compared with the results presented in the respective publications. A) When comparing between Job’s syndrome patients (HIES_control_0) and healthy controls (Healthy_control_0), the mean fold-change of CD151 was 1.7 on the GXB (GSE8507-PBMC). The reported value in Holland et al (2010) was 2.0 (38). B) When comparing between PBMC samples from egg allergic patients (Allergic_Egg) and allergic egg-tolerant controls (Allergic_Control), the mean fold-change of CEACAM1 was 1.69 on the GXB (GSE88796). Kosoy et al. (2016) reported a mean fold-change of 1.6 for the same gene (39). The overall trends of gene expression are conserved between GXB and published data.

## DISCUSSION

### Potential application of the dataset collection

To demonstrate the potential use of the dataset collection, gene expression profile of different tissues for house dust mite (HDM) allergy were compared. Nasal epithelial (GSE19190) and PBMC-derived CD4 T cells (GSE70760) gene expression profiles (detailed in Table 2) were compared. In both dataset, genes associated with the Th2 pathway/axis were increased in HDM-sensitized patients compared against healthy controls (32,33); illustrating that Th2 response may results in the symptomatic phase. This is further supported by the IgE levels and sIgG/sIgE ratio in the same study (33).

SERPINB2, a gene coding for the inhibitor of plasminogen activator PAI-2, is markedly upregulated in patients in both datasets, hence tissues (Suppl. Figure 1A from GSE70760 and Figure 1B from GSE19190). This is consistent with previous association of asthma severity and biomarker panel including SERPINB2 from PBMC (34) and correlation of SERPINB2 expression in respiratory epithelial cells with atopic asthma severity (35). SERPINB2 has been reported to have a role in the interleukin-12-mediated signaling pathway; evidence from mice showed that SERPINB2 regulates IFNg production, causing down regulation of Th1 cytokines in macrophages (36).

The expression profiles of SERPINB2 in both tissues suggest an important role of the gene as a first line regulator of immune response, perhaps by preventing excessive Th1 response. Interestingly, IFNg was not decreased in PBMC-derived T cells, but IFNGR1 was, suggesting that in T cells, SERPINB2 may exert its role on the expression of the receptor rather than on IFNg itself. In Schroder et al (2010), stimulation of SERPINB2 −/− cells with antiCD40/IFNg resulted in greater Th1 cytokine production, thus supporting the idea that SERPINB2 affects IFNGR (36).

However, certain genes are found to be differently expressed between nasal epithelium or PBMC-derived T cells. An explanation is that the activation of Th1 immune response may be different in the target tissue and peripheral circulation. For example, IL1B, a Th1 promoting cytokine is found to be increased in CD4+ T cells in allergic patients (Suppl. Figure 1C, from GSE70760), but in the nasal epithelium, the gene is not upregulated in patients with severe case of allergic rhinitis (“uncontrol asthma”) compared against healthy control (Suppl. Figure 1D, from GSE19190). Further hypothesis-generating comparisons can be made by comparing the top differently expressed genes in both datasets as listed in Suppl. Table 1.

### Conclusion

The dataset collection may be useful for exploring specific gene signatures in response to natural antigen/allergen exposure (i.e. allergic patients vs control) and delineate the major genetic drivers associated with increased level of IgE (i.e. cellular responses to specific antigen/allergen). Furthermore, comparative investigation of similarities and differences in expression of genes between datasets can highlight key mechanistic differences of immune signaling and/or provide insights for tissue-targeted intervention. In compiling the present dataset collection, we hope to offer a resource that may improve accessibility of public omics data to researchers in this field.

## DATA CITATION

1. IgE_GXB_Database *Figshare* http://doi.org/10.6084/m9.figshare.7176851 (2018).

## AUTHOR CONTRIBUTIONS

MG and SSYH contributed to <u>conceptualization</u>. MG, SSYH, and FA contributed to <u>data curation</u> and <u>validation</u>. MG and SSYH led, and FA supported, <u>investigation</u> and <u>visualization</u>. MG and SSYH performed <u>formal analyses</u>. SB and MT contributed to the maintenance of <u>software</u>. MG contributed <u>writing – original draft, methodology</u>, and <u>project administration</u>. MG and SSYH led, and FA, SB, MT, DC supported, <u>writing – review & editing</u>. MG lead and SSYH supported <u>supervision</u>. DC contributed <u>funding acquisition</u> and <u>resources</u>. The contributor’s roles listed above (underlined) follow the Contributor Roles Taxonomy (CRediT) described in Nature Communication 2014 (37) and managed by The Consortia Advancing Standards in Research Administration Information (CASRAI) (https://casrai.org/credit/).

## ACKNOWLEDGEMENT

We would like to thank all the investigators who decided to make their datasets publicly available by depositing them into the NCBI GEO repository.

## LIST OF SUPPLEMENT TABLES AND FIGURES

**Suppl. Table 1.**
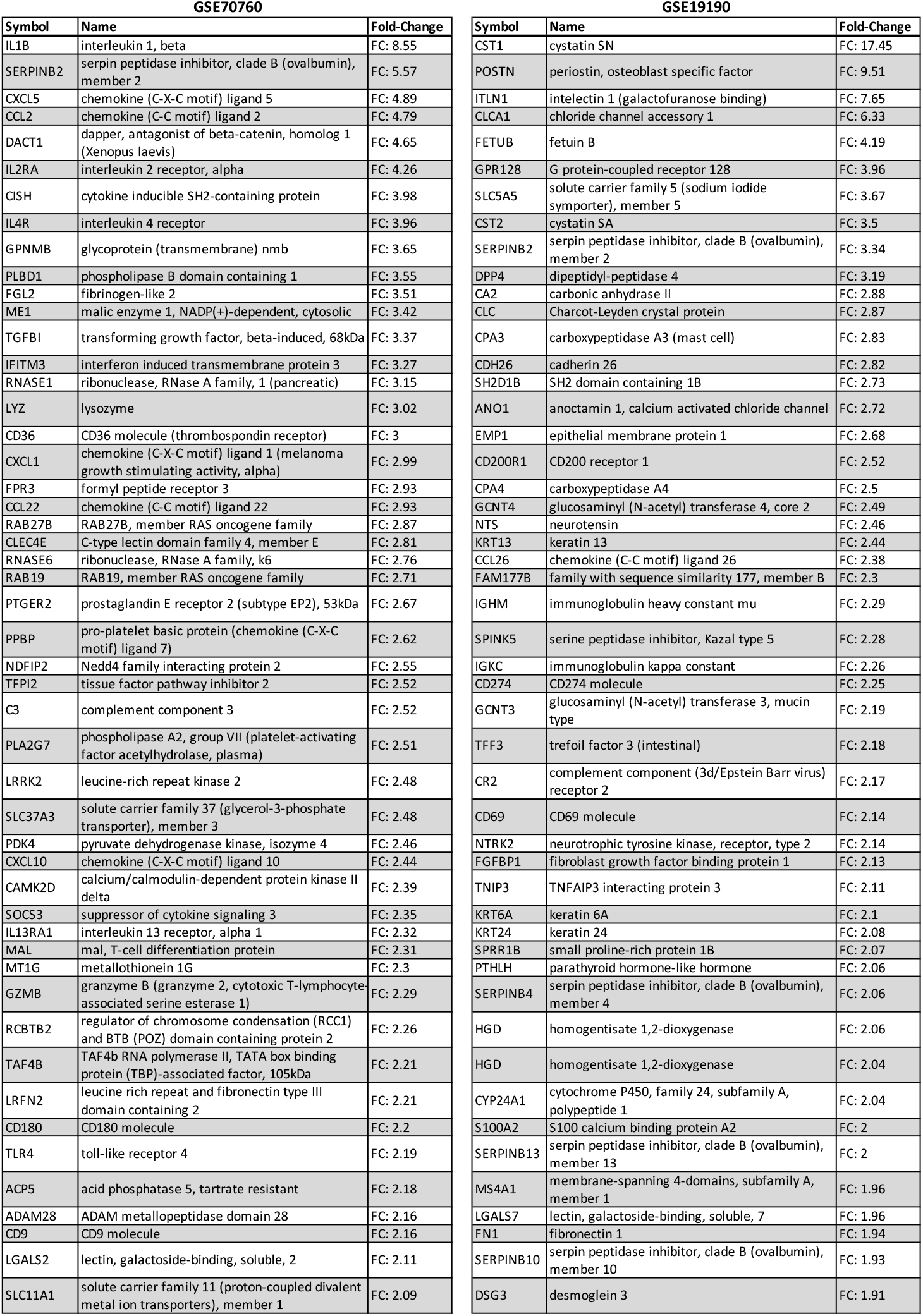
Top 50 differently expressed genes in datasets GSE70760 and GSE19190 as represented on GXB.

**Supplement Figure 1.**
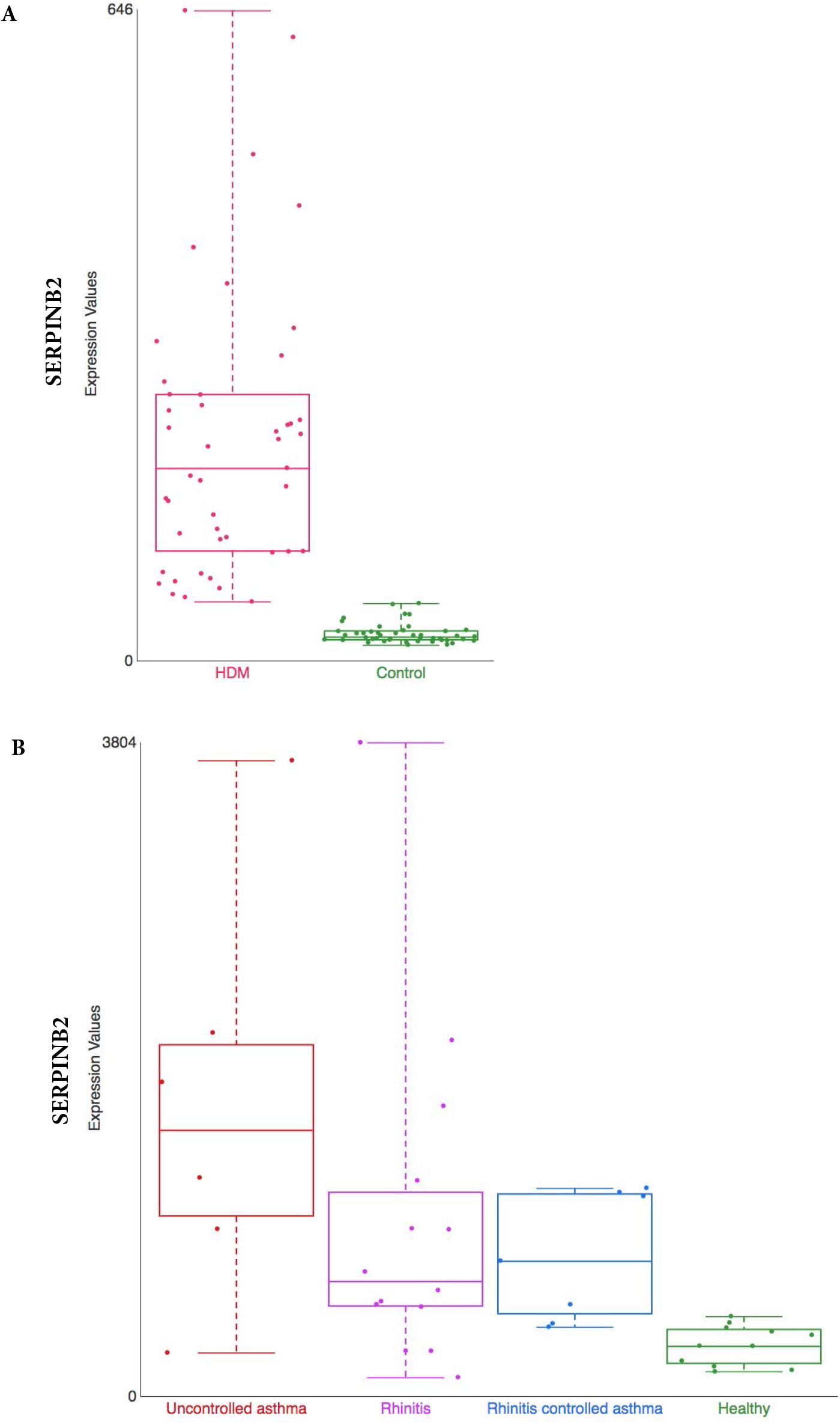
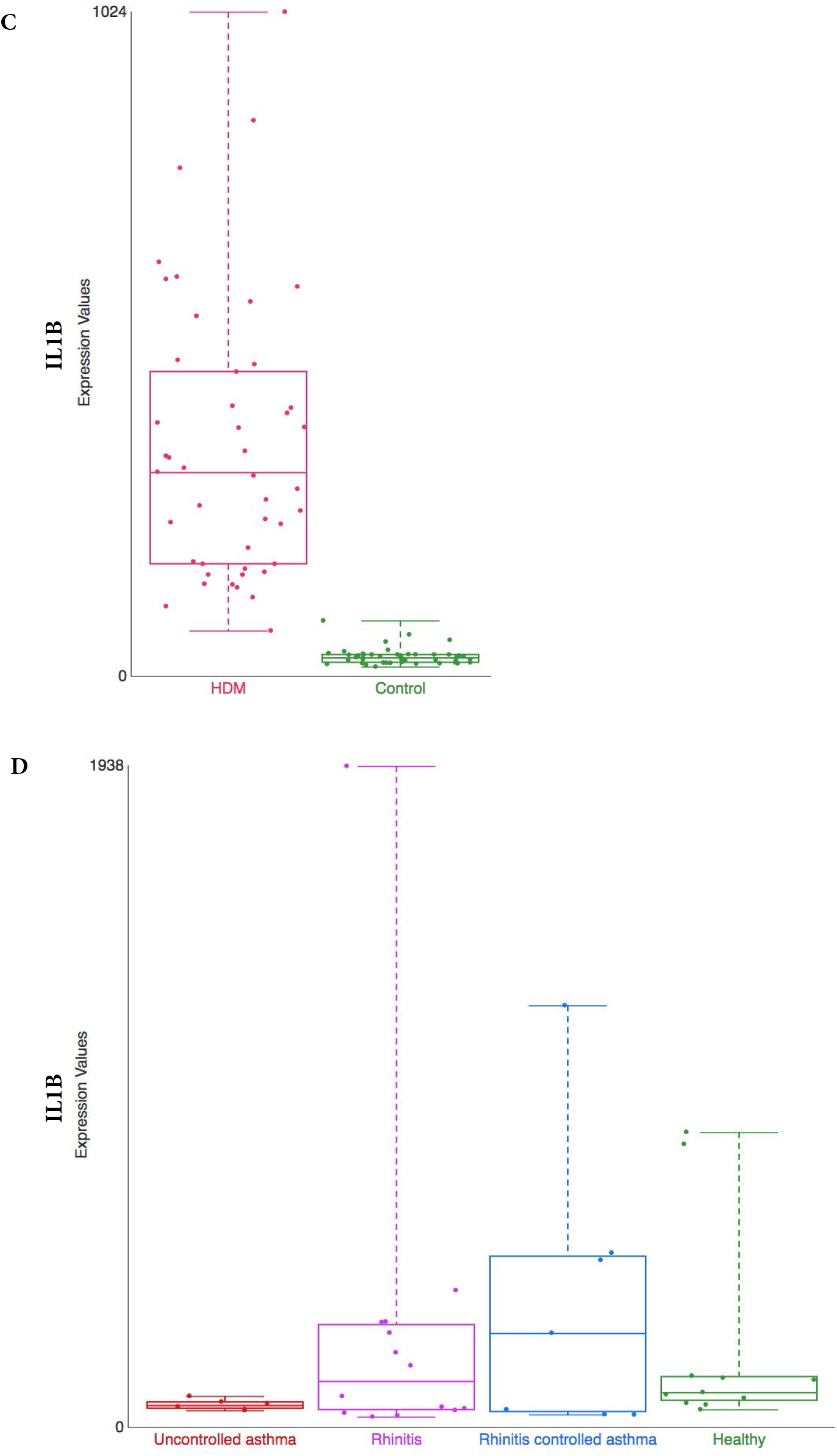
Comparison of gene expression profile in different tissue from house dust mite (HDM)-sensitized individuals. Gene expression data from two datasets present in our collection are shown: 1) Gene expression patterns in house dust mite stimulated CD4 T cells and IgG to IgE ratios - GSE70760, and 2) Distinct epithelial gene expression phenotypes in childhood respiratory allergy - GSE19190 - Disease State. SERPINB2 (A and B) and IL1β (C and D) gene expression. HDM = house dust mite; Control = Healthy = healthy individuals; AND Uncontrolled asthma = individuals with rhinitis and uncontrolled asthma (further definition can be found in the original study description (32).

**Supplement Figure 2.**
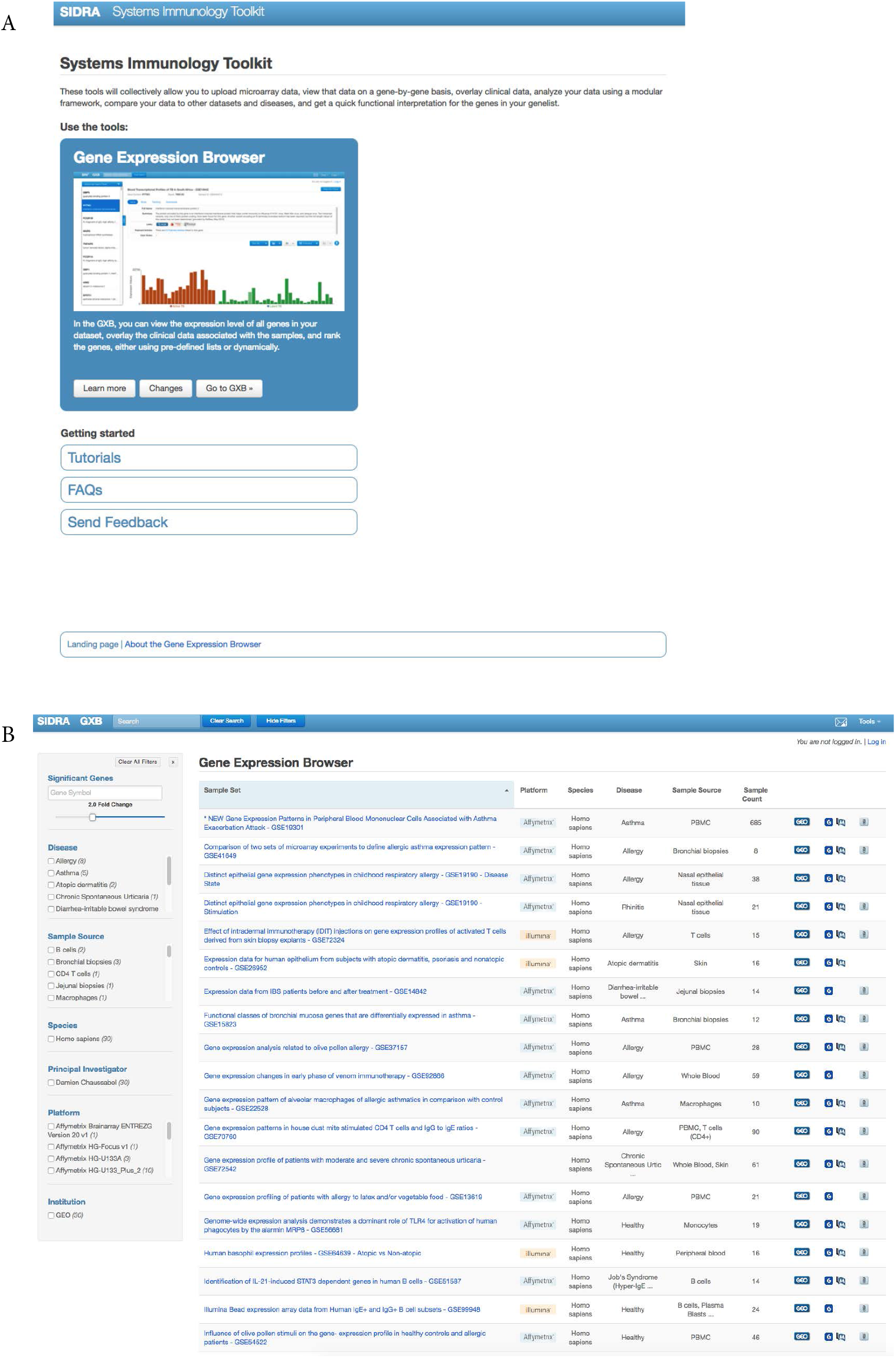
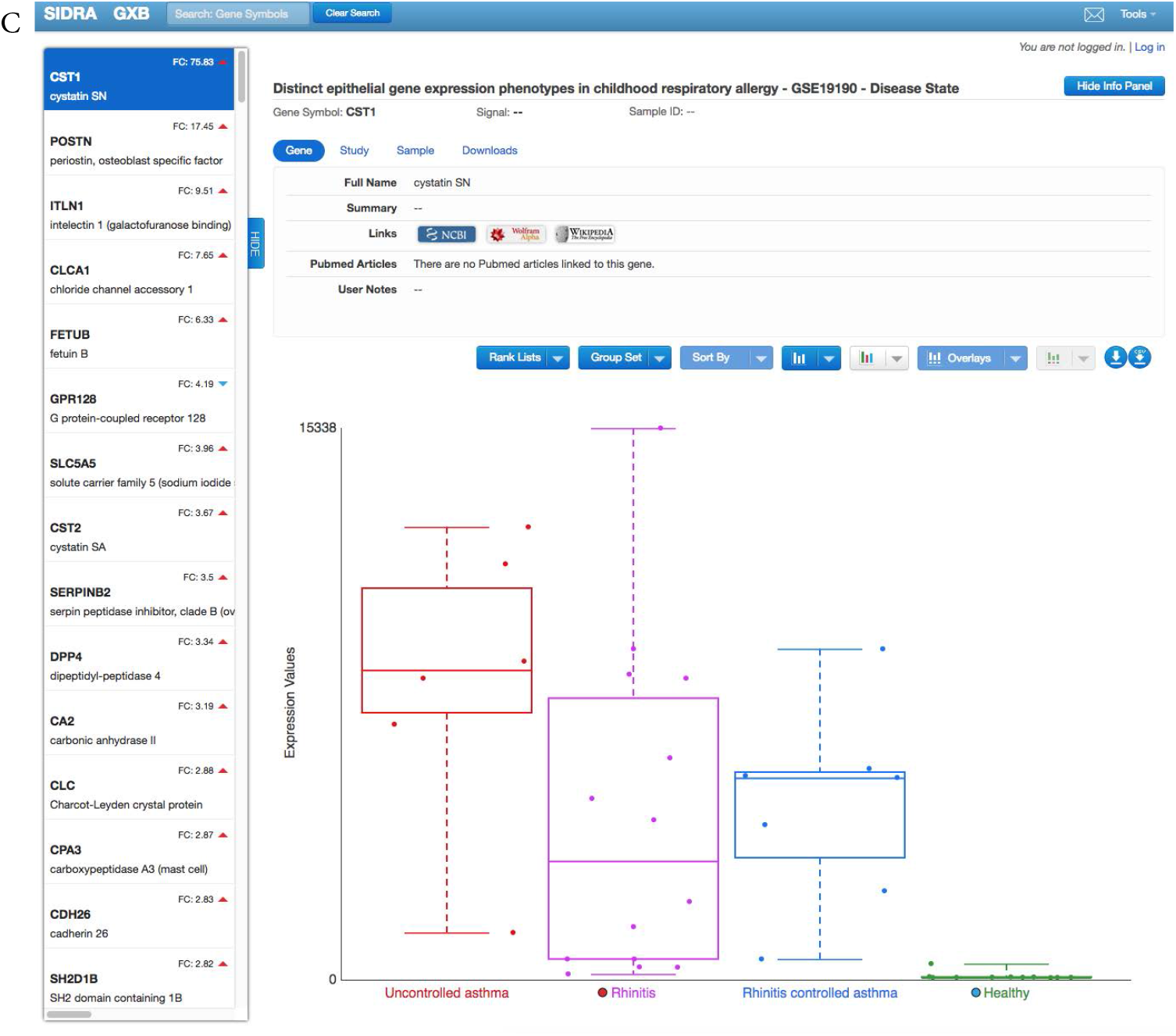
GXB interfaces. The end-users interact with 3 main interactive pages: the Landing Page (A), Dataset Browser (B) and Graphical (C) interfaces. A detailed description of the interface and functionalities have been previously described (PMID 26088622).

## SUPPLEMENTAL INFORMATION

**Supplement Information 1**. Calculation of FC expression.

GXB uses the geometric means of replicates of each sample and calculates FC via the difference (for data in log2 scale) or the ratio (for data in linear scale) of the means. The data displayed in GXB are linear scale FC. The following example illustrates the calculation:

**GSE ID**: GSE54336
**Type of file**: *.soft file (The data deposited by the contributor were analyzed with Partek Genomic Suite 6.6 using Affymetrix default analysis settings, quantile normalization and RMA background correction)
**Number of samples**: 6
**Number of groups**: 3 (G1TEPP, G1V, and Mock)
**Groups compared** (for this example): G1TEPP and G1V
**Gene symbol**: DUSP2
**Probe set ID**: 204794_at

### Sample Information

GSM1313408 A2EN cells_Chlamydia G1TEPP_4h_rep1

GSM1313409 A2EN cells_Chlamydia G1TEPP_4h_rep2

GSM1313410 A2EN cells_Chlamydia G1V_4h_rep1

GSM1313411 A2EN cells_Chlamydia G1V_4h_rep2

GSM1313412 A2EN cells_Mock infected_repl1

GSM1313413 A2EN cells_Mock infected_repl2

### Condition 1: G1TEPP (signal value in log_2_ scale)

GSM1313408: 6.31727

GSM1313409: 6.26554

Geometric Mean = sqrt (6.31727 * 6.26554)

➔ Geometric Mean = 6.29135183214

### Condition 2: G1V (signal value in log_2_ scale)

GSM1313410: 6.93182

GSM1313411: 6.88052

Geometric Mean = sqrt (6.93182 * 6.88052)

➔ Geometric Mean = 6.90612236689

### Calculation of FC expression (G1TEPP/G1V) and transformation to linear scale FC

log_2_ FC = Condition 1 - Condition 2

log_2_ FC = 6.29135183214 - 6.90612236689 = −0.61

linear FC = Antilog (−0.61) = 2^(−0.61)^ = 0.65

Mathematical transformation when linear FC is less than 1: −1/(FC)

➔ −1/0.65 = −1.52 (i.e. down regulation in linear scale)

